# Minimal clustering and species delimitation based on multi-locus alignments vs SNPs: the case of the *Seriphium plumosum* L. complex (Gnaphalieae: Asteraceae)

**DOI:** 10.1101/2021.03.21.436318

**Authors:** Zaynab Shaik, Nicola G Bergh, Bengt Oxelman, G Anthony Verboom

## Abstract

We applied species delimitation methods based on the Multi-Species Coalescent (MSC) model to 500+ loci derived from genotyping-by-sequencing on the South African *Seriphium plumosum* (Asteraceae) species complex. The loci were represented either as multiple sequence alignments or single nucleotide polymorphisms (SNPs), and analysed by the STACEY and Bayes Factor Delimitation (BFD)/SNAPP methods, respectively. Both methods supported species taxonomies where virtually all of the 32 sampled individuals, each representing its own geographical population, were identified as separate species. Computational efforts required to achieve adequate mixing of MCMC chains were considerable, and the species/minimal cluster trees identified similar strongly supported clades in replicate runs. The resolution was, however, higher in the STACEY trees than in the SNAPP trees, which is consistent with the higher information content of full sequences. The computational efficiency, measured as effective sample sizes of likelihood and posterior estimates per time unit, was consistently higher for STACEY. A random subset of 56 alignments had similar resolution to the 524-locus SNP data set. The STRUCTURE-like sparse Non-negative Matrix Factorisation (sNMF) method was applied to six individuals from each of 48 geographical populations and 28023 SNPs. Significantly fewer (13) clusters were identified as optimal by this analysis compared to the MSC methods. The sNMF clusters correspond closely to clades consistently supported by MSC methods, and showed evidence of admixture, especially in the western Cape Floristic Region. We discuss the significance of these findings, and conclude that it is important to *a priori* consider the kind of species one wants to identify when using genome-scale data, the assumptions behind the parametric models applied, and the potential consequences of model violations may have.

## 1.1 Species delimitation using multi-locus DNA sequence data

Building on a long history of conceptual development (Darwin 1859; Dobzhansky 1937; Mayr 1942; de Queiroz 1998; Wilkins 2009; Wiley & Lieberman 2011), species delimitation has recently moved into the genomic sphere with the development of methods which use sequence data from multiple, putatively unlinked DNA loci to infer species boundaries in the context of the multi-species coalescent (MSC; O’Meara et al. 2006; Yang & Rannala 2010, 2014; Zhang et al. 2011; Leaché et al. 2014a; Jones et al. 2015; Yang 2015; Jones 2017; Rannala & Yang 2017). As currently implemented, these methods generally assume that species conform to the conditions of the Wright-Fisher model (Fisher 1930; Wright 1931; Hein et al. 2004), namely random mating, population size constancy, non-overlap of generations and no migration (Hein et al. 2004), with the consequence that the species they reveal can be viewed as population lineages (Darwin 1859) or minimal evolutionary species (de Queiroz 1998). The last decade has seen important advances in the development of species delimitation methods based on DNA sequence data, as applied to both single-locus (e.g., GMYC: Pons et al. 2006; Fujisawa & Barraclough 2013; PTP: Zhang et al. 2013) and multilocus data (SpedeSTEM: Ence & Carstens 2010; BP&P: Yang & Rannala 2010, 2014; Bayes Factor Delimitation: Aydin et al. 2014; Grummer et al. 2014; Leaché et al. 2014a; DISSECT/STACEY: Jones et al. 2015; Jones 2017). Most real applications of the MSC, however, address situations in which the pervasiveness of incomplete lineage sorting and other processes that confound species tree inference limit the power and usefulness of single-locus methods, while at the same time rendering the concatenation of multiple loci inappropriate (Felsenstein 2004). These circumstances require the application of multilocus methods and which are, accordingly, the focus of this paper.

Fujita et al. (2012) partitioned MSC-based delimitation methods into “verification” methods, which select among a set of predefined delimitation schemes, and “discovery” methods, which select among all possible delimitation schemes. Bayes Factor Delimitation (BFD) is a verification approach which evaluates a set of alternative classifications using marginal likelihood estimates obtained in an MSC model framework. It can be applied to biallelic marker data (Leaché et al. 2014a), such as single nucleotide polymorphisms (SNP) or amplified fragment length polymorphisms (AFLP), or to sequence data (Aydin et al. 2014; Grummer et al. 2014), with marginal likelihood estimates from SNAPP (Bryant et al. 2012) or *BEAST (Heled and Drummond 2010), respectively, being used to compare the fits of the data to alternative species delimitation schemes. By contrast, discovery methods, which include recent versions of BP&P (Yang & Rannala 2014; Rannala & Yang 2017) and DISSECT (Division of Individuals into Species using Sequences and Epsilon-Collapsed Trees; Jones et al. 2015) as implemented (with improvements) in STACEY (Species Tree and Classification Estimation, Yarely; Jones 2017), use aligned sequence data to simultaneously derive posterior estimates of the gene trees, species delimitation, species tree and other model parameter distributions. As such they require no *a priori* specification of candidate delimitation schemes. Although Bayesian verification and discovery implementations differ in approach, they are expected to retrieve identical or similar species (Barley et al. 2018) so long as (i) the globally optimal or near-optimal delimitation schemes are included in the set of schemes considered by the verification methods, and (ii) the Monte Carlo Markov Chains (MCMCs) are run to convergence for the posterior distributions of the parameters estimated. This is often considered difficult to achieve, and to our knowledge, the sequence alignment-based methods have not been applied to more than a handful of empirically sampled loci (Jones et al. 2015; Toprak et al. 2016; Wagner et al. 2017; Matos-Maravi et al. 2019). One might assume that BFD using SNAPP would allow examination of a greater number of loci than with alignment-based methods, because the former explores a limited number of delimitation hypotheses, and because SNAPP bypasses the integration of gene trees completely and is, as a result, computationally less complex (Bryant et al. 2012). This, however, remains to be empirically tested.

A different, non-phylogenetic, model-based approach, widely used in population genetics and based on the STRUCTURE method of Pritchard et al. (2000), is to partition multilocus allelic variation into clusters which conform, as closely as possible, to the assumptions of Hardy-Weinberg and linkage equilibrium (Pritchard et al. 2000). This approach is typically applied within taxonomic species to reveal population structure and to compute individual “ancestry coefficients” which can be interpreted as the probability that a sampled allele is sourced from a particular gene pool (Frichot et al. 2014). Many variants of the original STRUCTURE method have been developed, including ADMIXTURE (Tang et al. 2005; Alexander, et al. 2009), fastSTRUCTURE (Raj et al. 2014), Snapclust (Beugin et al. 2018) and sNMF (sparse Non-negative Matrix Factorisation; Frichot et al. 2014), the last of which relaxes the assumption of Hardy-Weinberg equilibrium (Frichot et al. 2014). Of these, the latter three have runtimes more than an order of magnitude shorter than those of the STRUCTURE and ADMIXTURE (Frichot et al. 2014; Beugin et al. 2018), facilitating the computation of ancestry coefficients for large molecular data sets comprising many loci and/or many accessions. Although these genetic clustering algorithms resemble MSC-based species delimitation approaches in that clusters of objects (in this case, alleles) are optimized within a probabilistic framework, they differ in that genetic structure is modelled in a way that fails to consider the degree of relationship among populations and the pattern of historical population divergence. Thus, although they offer several benefits, including flexibility in the types of data which they can use, relatively short runtimes and a long history of application, they may yield clusters which do not correspond closely to the branching pattern of the species tree (Kalinowski 2011). Nonetheless, these approaches are commonly used as a starting point for species delimitation studies, with Carstens et al. (2013) recommending their application in conjunction with other methods whose assumptions may be violated by the system under study.

### 1.2 The <u>Seriphium plumosum</u> complex

This paper assesses the utility of two fully parameterised MSC-based clustering methods and a genetic clustering algorithm for the purpose of delimiting species within a widespread and morphologically variable lineage of Cape plants. Since its initial description (Linnaeus 1753), between one and four species have been recognised within *Seriphium plumosum* L. (*Seriphium* hereafter abbreviated as *S*.; Thunberg 1800; Lessing 1832; Candolle 1838; Harvey 1865; Levyns 1937; Koekemoer 2016). In a recent taxonomic treatment, Koekemoer (2016) segregated *Seriphium* from the genus *Stoebe* (hereafter abbreviated as *St*.), and at the same time using a non-statistical, morphology-based approach merged three formerly recognised species, *St. plumosa* (L.) Thunb., *St. burchellii* Levyns and *St. vulgaris* Levyns into a single species under the name *S. plumosum*. The result is a much-expanded concept of *S. plumosum*, comprising an assemblage of plants showing a diversity of growth forms, synflorescence and leaf arrangements, and leaf indumentum structures. The morphological variation shown by the complex, although subtle, is paralleled by the diversity of edaphic and climatic conditions it occupies; *S. plumosum* associates with winter, aseasonal and summer precipitation regimes (Supplementary Data), an elevational range from sea level to >1900 m, and a range of lithologies from quarzitic and shale-based substrates to calcareous and unconsolidated dune along the coast. Apart from its ecological and morphological diversity, the genetic integrity of *S. plumosum* is brought into question by the sheer extent of its distribution range (Supplementary Data), which considerably exceeds that of its immediate relatives (Koekemoer 2016) and indeed of most Cape plants (Latimer et al. 2005). The presence of multiple, independent, lineages within *S. plumosum* is further suggested by variation in the species’ flowering time across its range (Koekemoer, 2016), particularly in view of the potential efficacy of flowering time shifts in interrupting between-population gene flow (Devaux and Lande 2009).

### 1.3 Aims

The central aim of the present study is to assess the performance and behaviour of MSC-based and STRUCTURE-like delimitation methods when used with massively-parallel sequenced genome-scale molecular for species delimitation in genetically poorly-characterised species complexes such as *S. plumosum*. We examine the STRUCTURE-like approach sNMF (Frichot et al. 2014) here applied to biallelic SNP markers, as well as two recently-developed MSC-based methods: the BFD validation method (Leaché et al. 2014a), here applied to biallelic SNP markers, and the DISSECT/STACEY discovery method (Jones et al. 2015; Jones 2017) applied with sequence data (hereafter we refer to DISSECT/STACEY as STACEY). We examine the following questions and make the accompanying predictions:

i. What do potential incongruities between the clusters recovered by the discovery methods sNMF and STACEY suggest about the applicability of either approach for resolving species? Delimitation methods implementing the MSC resolve minimal clusters of *individuals* by optimizing fit to the genealogies predicted under the coalescent model (Kingman, 1982), while STRUCTURE-like algorithms cluster *alleles* by optimizing fit to linkage equilibrium. Differences in the input molecular data format (i.e., sequences verses SNPs) and the divergent conceptual approaches of either approach suggest cluster incongruities are to be expected, and can possibly inform the extent to which model assumptions are met.
ii. How do computation speeds and the clusters recovered by BFD and STACEY compare when using genome-scale data? Empirical applications of BFD typically include a fraction of the total number of species delimitation schemes possible, and BFD sensu Leaché et al. (2014a) uses biallelic markers rather than full sequences. In terms of computational tractability, we expect BFD will emerge as preferred over STACEY, and as BFD and STACEY are both implementations of the MSC, we expect they should provide support for the same clusters.
iii. Given the insights provided by (i) and (ii), how can molecular data on the scale captured using massively-parallel sequencing platforms be most appropriately applied for informing revised taxonomies?

## 2 Materials & methods

### 2.1 Sampling

Populations representing all previously recognised species in the *S. plumosum* L. complex and representing its full geographic range in South Africa were sampled between April 2017 and April 2018 from 47 localities (Supplementary Data). A single population of *S. cinereum*, situated near Cape Town, was also sampled (Supplementary Data) since a phylogenetic analysis of ITS and plastid Sanger sequences (N.G. Bergh, personal communication) has identified this as being the putative sister species of *S. plumosum*. In total, 48 localities were sampled; 47 for *S. plumosum* and one for *S. cinereum* (Supplementary Data). Since all sampled plants had very small (1-5 mm long) leaves, short (ca. 5 cm) cuttings were collected and stored on silica-gel for DNA isolation. Six individuals were collected at each locality for a total of 288 samples. Collected plants were spaced ≥ 5m apart to reduce the chance of sampling close relatives. A single pressed voucher specimen for each population has been deposited at the Compton Herbarium, Kirstenbosch, South Africa (NBG; Supplementary Data).

### 2.2 Data generation

#### Laboratory procedures

Since genotyping-by-sequencing (GBS) requires high-quality DNA, extraction was performed within one week of collection to minimize degradation. For each sample, 100 µg of silica-dried leaf material was pulverised in liquid nitrogen prior to DNA extraction using the DNEasy plant mini kit (*Qiagen, Venlo, Netherlands*) following the manufacturer’s instructions but with final elution in 30 μl TE buffer. While this protocol generally yielded samples having concentrations ≥50 ng/μl and 260/280 absorbance ratios between 1.8 and 2, a few samples required further concentration by speed-vacuuming for 10 minutes at 43°C using the Savant SpeedVac concentrator SC210a (*ThermoFisher Scientific, Waltham, MA USA*). Extracts were stored at -80 °C, before being express-shipped to Novogene Genome Sequencing Company Ltd. (Beijing, China) for GBS. Here, sample genomes were fragmented using MseI, HaeIII, and MspI, and the resulting fragments ligated with adaptors. Adaptor-ligated fragments about 350 base-pairs (bp) long according to the Agilent 2100 bioanalyzer (*Agilent, California, United States*) were recovered using size selection, amplified by quantitative real-time PCR (qPCR), and paired-end sequenced on the Illumina HiSeq PE150 (*Illumina, Inc*.) system.

#### Data filtering, sequence assembly, variant calling

Raw sequence data were filtered to exclude paired reads with adaptors still attached, >10% ambiguously called nucleotides, and paired reads where >50% of either read comprised low quality nucleotides (quality score ≤Q5). The full set of filtered reads was used to assemble a *de novo* reference genome using SOAPdenovo version 2.04 (Luo et al. 2012) with parameter settings [-K 41 -R -d 2 -p 10] and minimum quality thresholds as follows: contigs (consensus sequences) N50 ≥ 30 kilobase pairs and scaffolds N50 ≥ 1 mega bp. After sequencing, *de novo* reference genome assembly, and preliminary filtering by Novogene as detailed above (Novogene Genome Sequencing Company Ltd. 2016), we generated a single concatenated variant data set across all 288 accessions by quality-trimming and mapping reads to the *de novo* reference genome, calling variant sites, and filtering the variant sites using Puritz et al.’s (2014) dDocent pipeline and R version 3.5.1. (R Core Development Team 2018). Quality trimming of cleaned reads was performed using Trimmomatic (Bolger et al. 2014), which removed any remaining adaptor contamination and low-quality bases (<Q20, or error rate 1% error rate) from the beginnings and ends of reads. Trimmed reads were mapped to the *de novo* reference genome using the MEM algorithm of the Burrows-Wheeler Aligner (BWA), the preferred algorithm for low-divergent sequences of greater than 70 bp long (Li & Durbin 2010), with default conservative mapping parameter settings [-A 1 -B 4 -O 6]. Parallel variant calling was performed across 64 processing cores with 128 GB of memory (University of Cape Town ICTS High Performance Computing Centre) using FreeBayes version 1.2.0 (Garrison & Marth 2012), and a complete variant call file (VCF) created using VCFtools version 3.0 (Danecek et al. 2011). Genotyped variants were advance-filtered as part of the dDocent pipeline using VCFtools to produce a final SNP data set, with population-specific genotype call rate set to 10% across five populations (Puritz et al. 2014). A subset of the resulting VCF comprising 36 accessions with no missing data across loci (VCF-36 hereafter) was generated in R using the package *vcfR* version 1.8.0 (Knaus & Grünwald, 2017). For BFD we used a 524-SNP subset of VCF-36 in which only one SNP per sequence alignment was retained across 36 accessions (Bryant et al., 2012).

#### Assembly of sequences for multiple alignments

Sequence alignments for STACEY analysis were assembled from the GBS data set by (i) querying the identity of the unique scaffold to which each short read containing a SNP in VCF-36 was aligned, (ii) using the scaffold identifiers to subset filtered reads in the binary alignment/map (BAM) files (generated by mapping above), and (iii) using the filtered BAM reads to assemble two haplotype sequences per individual at each locus. Haplotyping is an efficient method for detecting multicopy loci and reducing nucleotide ambiguities (Zhi et al. 2012; Willis et al. 2017; Andermann et al. 2019), and for this reason haplotypes are preferred over consensus sequences when using STACEY (Andermann et al. 2019). Given their large size, BAM files were individually loaded into R using the package *ssviz* version 1.16.1 (Low 2019) and filtered to exclude reads not containing variants present in VCF-36. Reads aligned to the same scaffold identifier were aligned using the R package *muscle* version 3.26.0 (Edgar 2004). Aligned reads obtained from the same individual were then haplotyped using the R package *pegas* version 0.12 (Paradis 2010) and haplotypes ordered by descending read depth (number of reads supporting a haplotype). To exclude multicopy loci, alignments with third and fourth haplotypes of read depth ≥3 were discarded, based on the assumption that all individuals are diploid (Semple & Watanabe 2009; Vallés et al. 2013), while loci with either one or two haplotypes of read depth ≥3 (Andermann et al. 2019) were retained as orthologous. Paired haplotypes for each individual were then sorted by scaffold identifier, producing alignments containing up to 72 sequences (2 × 36 accessions). Where both haplotypes for an accession were absent at a locus on account of having >2 haplotypes of depth ≥3, or where only one haplotype of depth ≥3 was captured (either because of homozygosity or failed capture of a second haplotype), blank sequences matching the alignment length were inserted, but mean coverage was high (68.92/72) and there were few or no missing sequences per alignment. To further exclude non-orthologues, the absolute number of parsimony-informative sites across all samples was calculated for each locus using the R package *ips* version 0.0.11 (Heibl 2008), and loci containing >3 times the median number (n = 6 for our data) of parsimony-informative sites discarded. Despite the conservative default settings used when mapping reads to the *de novo* reference using the MEM algorithm, apparent multicopy loci (highly divergent haplotypes within individuals) remained in the resulting alignments. These were examined further by generating UPGMA trees (Sokal and Michener 1958) for each locus in the R package *phangorn* version 2.5.5 (Schliep 2011) and discarding as putative non-orthologues those loci in which ≥3 individuals had had haplotypes that were separated across the root node. All file conversions in the R environment were performed using the R package *Biostrings* version 2.50.2 (Pagès et al. 2019), and final alignments converted to FASTA format using the R package *seqinr* version 3.6.1 (Charif & Lobry 2007). Bioinformatic assembly of sequence alignments was performed in the R environment on a Lenovo X260 2.3GHz 4-core computer with 8 GB RAM.

### 2.3 Analysis

#### Gene pool assignment

The sNMF algorithm was applied to the broadest SNP data set comprising 28023 SNPs across the 288 accessions as implemented in the R package *LEA* version 2.2.0 (Frichot & François 2015). For each value of k between k = 2 and k = 20, 100 ancestry coefficient matrix replicates were computed (see Supplementary Data), where k represents the number of gene pools. The optimal number of gene pools (the value of k that, according to the least-squares criterion, best explains the genotypic data) was selected using the entropy criterion computed as part of sNMF for each k value as recommended by Frichot and François (2015). The Q matrices containing ancestry coefficient estimates generated by sNMF for the best-supported value of k were summarised using Clumpak (cluster Markov packager across K; Kopelman et al. 2015) which produces major and minor solutions to the least-squares minimisation algorithm based on the number of replicates supporting the solution for each value of k.

#### STACEY

Multiple haplotype alignments generated as described above were input into STACEY analysis (Jones, 2017), a method based on a modified birth/death model (Jones et al. 2015) under which divergence times associated with very recent splits are approximated as zero. This allows for the identification of minimal clusters of individuals (e.g., single organisms) which correspond to populations (“species”) as modelled in the MSC model (Heled & Drummond 2010; Jones 2017). The STACEY template in BEAUTi version 2.5.0, Java version 1.8.0_91 (Bouckaert et al. 2014) which is used to assemble XML files for BEAST analysis failed when hundreds of loci were imported. The XML file for STACEY input was thus partly assembled in BEAUTi using the STACEY template, with the remainder of the assembly automated in R using a custom script available on GitHub (https://github.com/zaynabthebotanist). Each accession was defined *a priori* as a minimal cluster, except in population number 32 (*S. cinereum*) where three accessions sampled from the same sampling locality were defined as a minimal cluster. Since each accession was represented by two haplotypes (one maternal and one paternal), the number of sequences per locus and minimal cluster ranged from two to six.

The analysis was implemented in BEAST 2 version 2.6.0 with STACEY version 1.2.5 (Bouckaert et al. 2014). A Jukes-Cantor (Jukes & Cantor 1969) substitution model and a strict molecular clock were used, with the substitution rate of the first (arbitrarily selected) partition set to 1, and those of the remaining partitions estimated relative to this. Collapse height (ε) was set to 10^-5^ (Jones et al. 2015). Priors on the clock rates were all drawn from independent lognormal distributions with (log) mean and (log) standard deviation set to 0 and 1.25, respectively; hereafter written as: lognormal (0, 1.25). The growth rate prior was drawn from a lognormal (5, 2) distribution. The population size prior was lognormal (-7, 2), and death rate and collapse weights were uniformly distributed using a β(1,1) distribution, implying they were uninformative. We ran six parallel MCMC analyses for several billion generations with 32 GB per CPU allocated to run on resources provided by UNINETT Sigma2, the National Infrastructure for High Performance Computing and Data Storage in Norway (Uninett Sigma2 2020) on the “Saga” cluster, sampling parameters and trees every 500 000^th^ generation. Because preliminary results indicated that the root node depths of some gene trees were exceptionally high compared to the species tree (by one to two orders of magnitude), the alignments were inspected manually. We identified stretches of poorly aligned sequences in 24 of the 524 alignments, usually as extra flanking, but strongly conflicting sequences. This analysis is labelled 1 in Table 1. For further analyses (versions 2a-2d, Table 1), these sections of sequence were replaced with question-marks in three of the runs and the MCMC repeated with the parameters and priors as the other analysis. Since all these analyses failed to converge and mix sufficiently, with effective sample sizes (ESS) for the posterior, likelihood, smcCoalescent, and TreeHeight estimates falling in the range 50-100 depending on burn-in, we also performed a less parameter-rich analysis on the curated alignments, in which the clock rates were all set to 1 (version 3, Table 1). Three runs (versions 0a-c, Table 1) were performed with gene-specific clock rates, and 56 randomly chosen alignments. SpeciesDelimitationAnalyser version 1.8 (Jones 2019b) was used to summarise the posterior frequencies of clusters based on the posterior sample of trees with burn-in determined by maximising the ESS values, ε = 10^-5^ and similarity cut- off set to 1. TreeAnnotator was used, with the same burn-in fraction, to summarize the maximum clade credibility (MCC) tree.

#### BFD analysis

Since the method BFD compares the marginal likelihood estimates of alternative species delimitations using Bayes Factors, we formulate alternative species delimitations for BFD *a priori* using past taxonomic treatments (Levyns 1937; Koekemoer 2016) as well as alternative groupings revealed by sNMF. We also calculate a marginal likelihood estimate for the Species/Minimal Clusters (SMCs) estimated by STACEY to assess agreement between STACEY and BFD. The least species-rich hypothesis is based on the most recently proposed taxonomy (Koekemoer 2016) which considers *S. plumosum* to be a single species. A second taxonomy-informed hypothesis is based on Levyns’ (1937) three-species taxonomy, which recognised under the generic epithet *Stoebe* L. *St. plumosa* sensu Levyns, a more narrowly circumscribed concept than *S. plumosum* sensu Koekemoer which is centred in the Cape Floristic Region (CFR) and extends into Zimbabwe, Namibia and Botswana; *St. burchellii* Levyns, which is confined to the Little Karoo region of South Africa; and *St. vulgaris* Levyns, which was applied to all the summer rainfall populations of *S. plumosum* sensu Koekemoer.

Marginal likelihood and species tree estimation using biallelic SNPs were implemented using SNAPP version 1.5.0 (Bryant et al. 2012) in BEAST 2 version 2.6.0 (Bouckaert et al. 2014). All BFD model runs included sequence data for the three individuals of *S. cinereum*. Single nucleotide polymorphism data in VCF format were converted to NEXUS format using PGDSpider version 2.1.1.3 (Lischer & Excoffier 2012). For each competing species delimitation hypothesis, XML files were generated in BEAUTi version 2.5.0 Java version 1.8.0_91 (Bouckaert et al. 2014). Marginal likelihood estimation was performed with stepping-stone analysis using the BEAST 2 application PathSampleAnalyser with 20 steps, each consisting of 3 × 10^5^ generations, and sampling parameters every 1000 steps. The alpha value was set to 0.3 (Leaché & Ogilvie 2016; Noguerales et al. 2018), while mutation rates u and v were both set to 1.0. A conservative set of delimitation priors (that favour lumping over splitting of clusters) as recommended by Leaché and Ogilvie (2016) were set for the expected genetic divergence prior Θ ∼ G(shape = 1, rate = 250) and speciation rate prior λ ∼ G(shape = 2, scale = 200), with means 0.004 and 400 respectively (See Leaché & Ogilvie [2016] for prior mean calculations). Marginal likelihoods for the taxonomy-informed hypotheses were estimated on an i7 3.6 GHz desktop computer with 16 GB of RAM and 8 CPUs. Marginal likelihoods for the remainder of the hypotheses were calculated using Linköping University’s National Supercomputer Centre’s “Tetralith” cluster (National Supercomputer Centre 2020) on a “fat” node with 384 GB of RAM and 20 CPUs (one CPU per path sampling step, steps run in parallel). To assess the consistency of marginal likelihood estimates, we ran five replicates for the delimitation hypothesis “sNMF1” (Table 2). Relative support for competing species delimitation hypothesis was evaluated using Bayes Factors (BF = 2 × [MLEmodel1 – MLEmodel2]) following the scale of Kass and Raftery (1995), in which BF > 10 is considered decisive support.

#### Species tree estimation

For the STACEY analyses and the BFD hypothesis with the greatest BF support, TreeAnnotator (version 2.6.0) was used to identify the topology with the greatest product of posterior branch support (the maximum clade credibility or MCC tree) from the posterior distribution of species trees (Leaché & Ogilvie 2016). Posterior node support was poor in the BFD MCC tree associated with the best-supported species delimitation hypothesis, suggesting failed convergence in the tree estimation SNAPP path sampling step of this analysis (Supplementary Data). Consequently, we ran a separate SNAPP tree estimation analysis using the same parameter settings as before for 2 × 10^6^ generations using Uninett Sigma2’s “Saga” cluster with 32 Gb of RAM on one CPU (as for the STACEY runs). FigTree version 1.4.3 (http://tree.bio.ed.ac.uk/software/figtree/) was used to visualise the MCC trees.

#### Computational runtimes

In order to compare the computational times for the STACEY and SNAPP runs relative to convergence (defined as effective sample sizes, ESS), we used the time per million generations estimated by BEAST and logged in the standard output. Due to run time restrictions on the computer cluster used, we only considered the output from the final 14-day run. Although this is a somewhat crude estimate, we consider it a useful measure of the computational intensity of the methods being compared.

## 3 Results

### 3.1 Sequence reads, SNP calling and alignment generation

Sequencing on the Illumina HiSeq PE150 platform yielded approximately 200,000 150 bp paired end reads per sample with an average read depth of 8, totalling more than 5.7 × 10^8^ raw sequence reads across all 288 accessions. This initial bioinformatic processing using the dDocent pipeline (read trimming, mapping, variant calling and filtering) produced 7 610 586 variant sites, including bi- and multi-allelic SNPs, indels, and composite indel and substitution events. After filtering reads using genotype depth, locus quality, minor allele frequency and genotype call depth, there remained 32 145 high-confidence variant sites across 288 individuals with 4.78% missing data. The 36-accession subset (VCF-36) generated from this data set and filtered to exclude variants with missing data contained 1809 biallelic SNPs.

The novel R-based bioinformatic workflow applied here produced 1329 sequence alignments. After filtering for the number of parsimony-informative sites, more than two significant haplotypes per individual, and haplotype separation across UPGMA tree root nodes, 524 alignments remained. These had a mean length of 155.43 bp (SD = 16.48) and contained a mean of 6.17 parsimony-informative sites (SD = 4.25), such that the full sequence alignment contained a total of 3 234 parsimony-informative sites across up to 72 haplotypes in 36 accessions.

### 3.2 sNMF analysis

Cross entropy criterion values were highly similar for k values between about ten and 17 (Supplementary Data), but achieved a minimum of 0.254 for k = 13 (Fig. 1e), suggesting the presence of around 13 putative gene pools; 12 in the *S. plumosum* complex (labelled A through L in Fig. 1a), and one corresponding to *S. cinereum* (Fig. 1a: M). The major mode for k = 13 (Fig. 1a), supported by 48 replicates, and the minor modes, supported by 23, 19 and ten replicates respectively (Fig. 1b-d) are largely consistent in the allocation of individuals to putatively independent gene pools or clusters, while all 100 sNMF replicates for k = 13 support the distinctness of *S. cinereum* (cluster M). Within the *S. plumosum* complex, clusters A – J house individuals distributed in winter- to aseasonal-precipitation zones, while K and L represent individuals from the summer-rainfall, eastern portions of South Africa. Clusters A – G are geographically confined to the Cape Fold Belt mountains: cluster A is a single population on a unique, high-altitude shale habitat; clusters B through F correspond to geographically-segregated populations of a common montane form, and cluster G corresponds to *St. burchellii* sense Levyns (1937). Individuals assigned to cluster G exhibit little allele sharing with other clusters, as do clusters I and J, and some samples in H. The clusters K and L, which exhibit low levels of gene pool mixing (particularly L; Fig. 1a-d), correspond to *St. vulgaris* sensu Levyns (1937).

**Figure 1.**
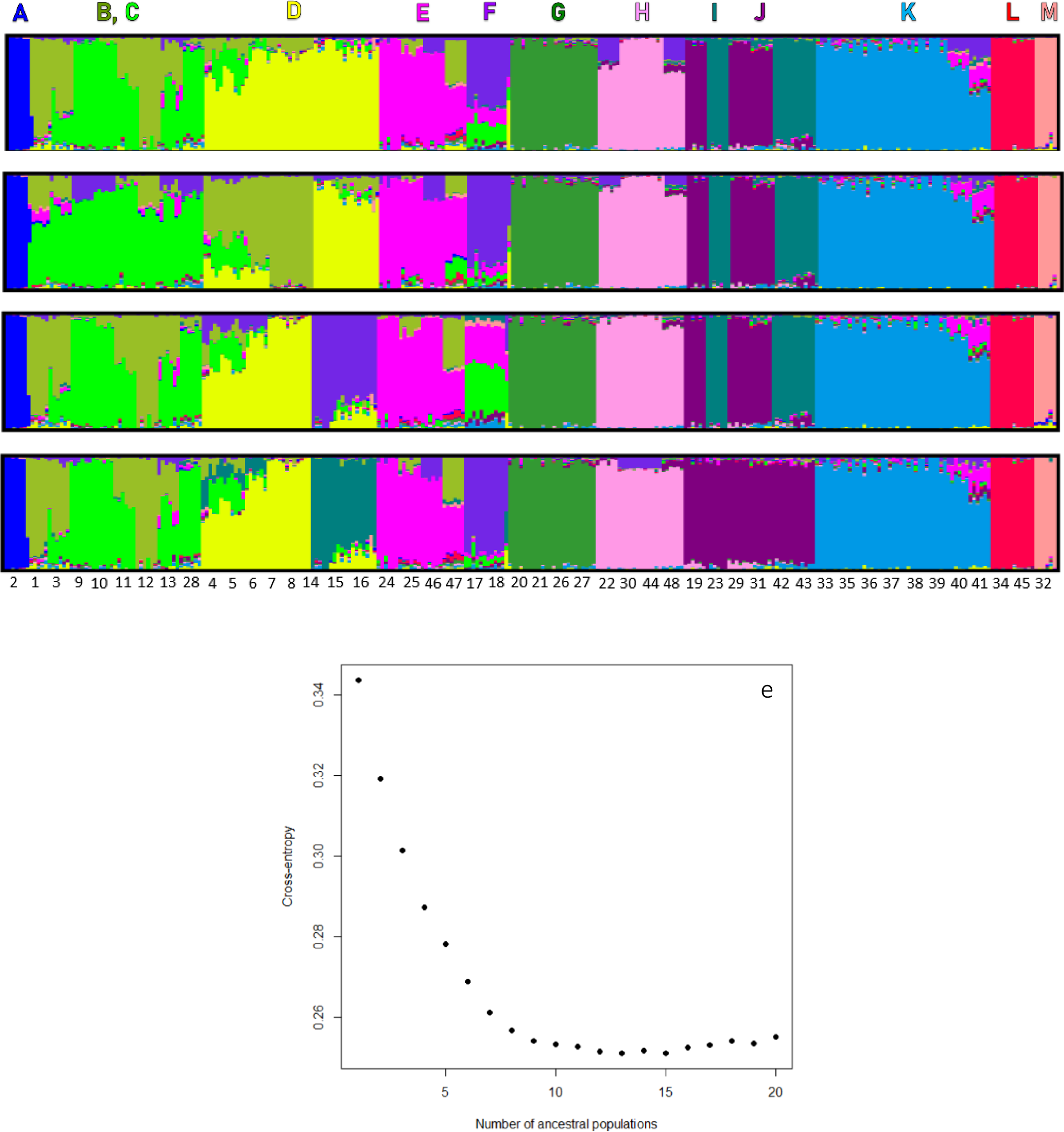
Plots of sNMF genomic assignment for the *S. plumosum* complex and *S. cinereum* for k = 13. Each individual is represented by a single vertical bar. Imputed gene pools (A-M) are indicated by colour. Major (a) and minor (b-d) cluster coefficient mode summaries are shown for all accessions (n = 288) from 48 numbered sampling localities. Letters above major mode correspond to the allele groupings supported under k =13. (e) The cross-entropy criterion for k = 2 though k = 20.

#### 3.3 STACEY analysis

Since convergence of the 524-locus analyses with each gene having its own clock rate was poor (versions 1-2d, Table 1), a less parameter-rich analysis with all clock rates equal was run for 5.688 × 10^9^ generations. With removal of the first 2.162 × 10^9^ generations as burn-in, all parameters of interest had ESS’s > 177 (Table 1). The minimal clusters and the associated tree resolved by this analysis and the six with individual clocks for the genes were similar (Supplementary Data) and, as such, we present only the results of the equal-clock rates analysis here (Fig. 2). Only one population pair (30 and 44, which are part of sNMF cluster H; Fig. 1a) was supported as conspecific, forming a minimal cluster in 75.32 % of the posterior samples (Fig. 2). Most of the population clusters identified by sNMF as distinct gene pools were resolved as monophyletic with posterior probability (PP) = 1.00 (Fig. 1a; Fig. 2). The clusters C and E (Fig. 1a) were not recovered as reciprocally monophyletic, but formed part of a larger well-supported (PP = 1.00) clade that also contained the clades B (Fig. 1a; Fig. 2) and F (Fig. 1a; Fig. 2), and the monophyletic cluster K (PP = 1.00; Fig. 1a; Fig. 2). The relationships portrayed in this tree are not strongly congruent with geography or rainfall seasonality, and the putative sister species *S. cinereum* (M) is more closely related to the remaining populations than is the unusual Cape Fold Belt population A.

**Figure 2:**
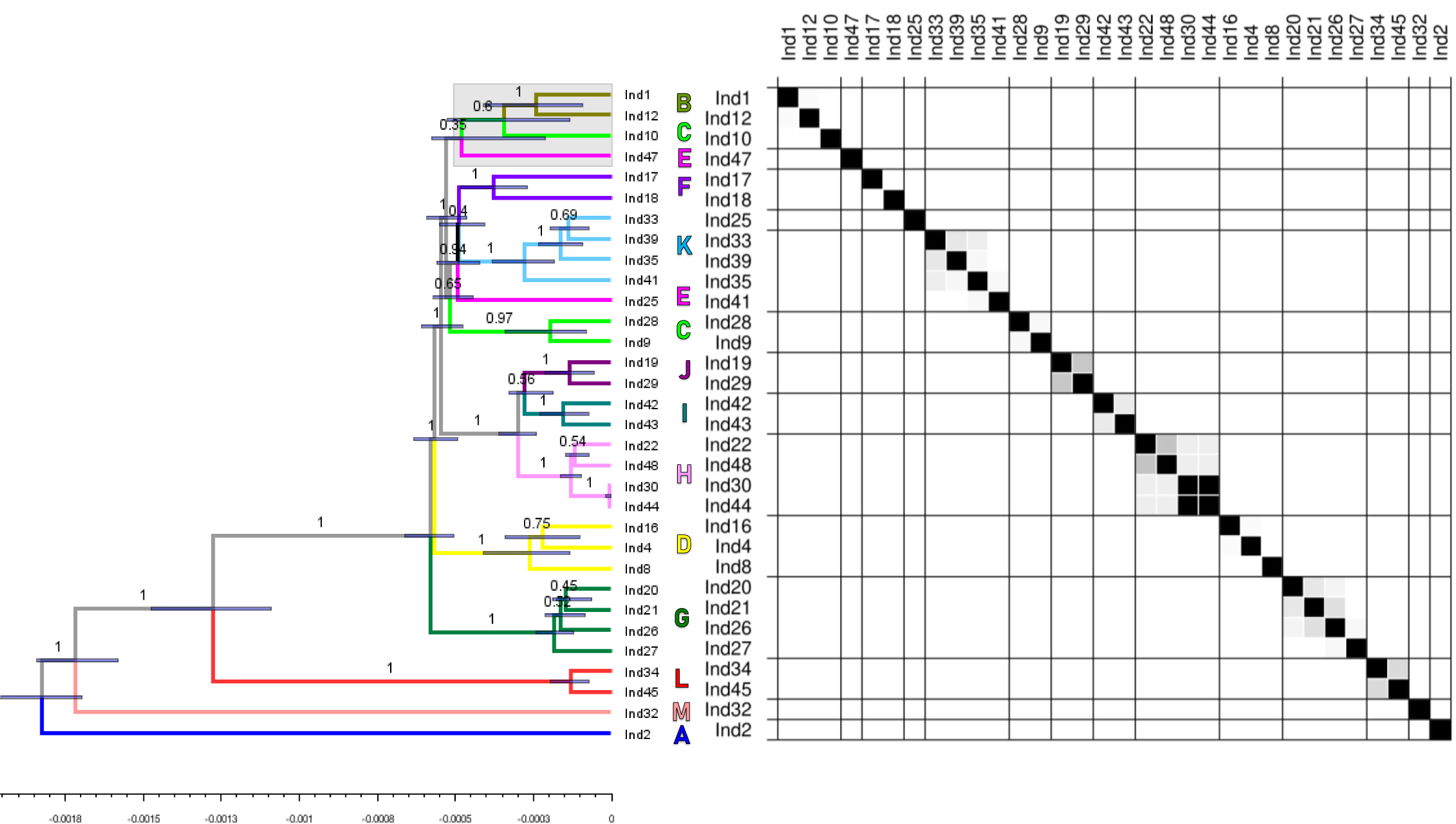
Species tree and two-way similarity matrix showing pairwise posterior probabilities of sampled individuals to belong to the same species based on STACEY analysis 3 of 524 sequence alignments for 36 accessions in *Seriphium cinereum* and the *S. plumosum* complex. Labels on tree branches denote posterior probabilities for clades of individuals, bars represent the 95% highest posterior densities for node heights. Tip accession labels and colours and letterings are consistent with those in Figure 1. The squares in the similarity matrix represent the posterior probability (white = 0, black = 1) that accessions belong to the same species or minimal cluster (sensu Jones et al., 2015) under ε = 10^-5^.

### 3.4 BFD of alternative species delimitation schemes

The clusters recovered by sNMF (Fig. 1a) provide support for several alternative delimitation hypotheses. Splitting or merging the coastal plain populations (Fig. 1a: I-J) and splitting or merging the admixed populations from the Cape Fold Belt mountains (Fig. 1a: B-F), excluding those populations corresponding to *St. burchellii* (Fig. 1a: G) produces four possible species hypotheses (Table 2). These four hypotheses, and those informed by prior taxonomies and by STACEY, test the division of the *S. plumosum* complex into between one and 30 hypothesized species.

The BFD analyses used the same 524 biallelic SNPs corresponding to the loci used in the STACEY analysis, and were run for 3 × 10^5^ generations for each of 20 path sampling steps. With removal of 3 × 10^4^ generations as burn-in, the average ESSs for the posterior and likelihood were 42.68 (sd = 38.43) and 31.80 (sd = 15.21) respectively. Bayes Factor comparison identified the species delimitation hypothesis based on the STACEY analysis as optimal (Table 2), the support for this hypothesis far exceeding that for any other. This was followed by the four sNMF-informed hypotheses which had rather similar BF support, and the three-species hypothesis based on Levyns’ (1937) taxonomy (Table 1). Bayes Factor support was lowest for the hypothesis based on current taxonomy, which treats *S. plumosum* as a single species (Table 2). The 3 × 10^5^-generation SNAPP species tree estimated in the BFD analysis with species boundaries informed by STACEY (Supplementary Data) supported the monophyly of the coastal dune populations (Fig. 1a: H), the Great Escarpment populations (Fig. 1a: L) and *St. burchellii* (Fig. 1a: G) as well-defined clades, but node support was frequently poor, as was node support for most of the remaining clades, which were inconsistent with both the sNMF results (Fig. 1a) and the STACEY MCC tree (Fig. 2). The two independent SNAPP MCC tree estimation analyses based on STACEY-informed clusters (Fig. 3) ran for more than 2 × 10^6^ generations each and appeared convergent topologically (Supplementary Data) but the TreeHeightLogger and likelihood failed to converge with ESSs of 24 and 81, respectively, in a combined trace with optimised burn-ins (10 and 50 percent respectively).

**Figure 3:**
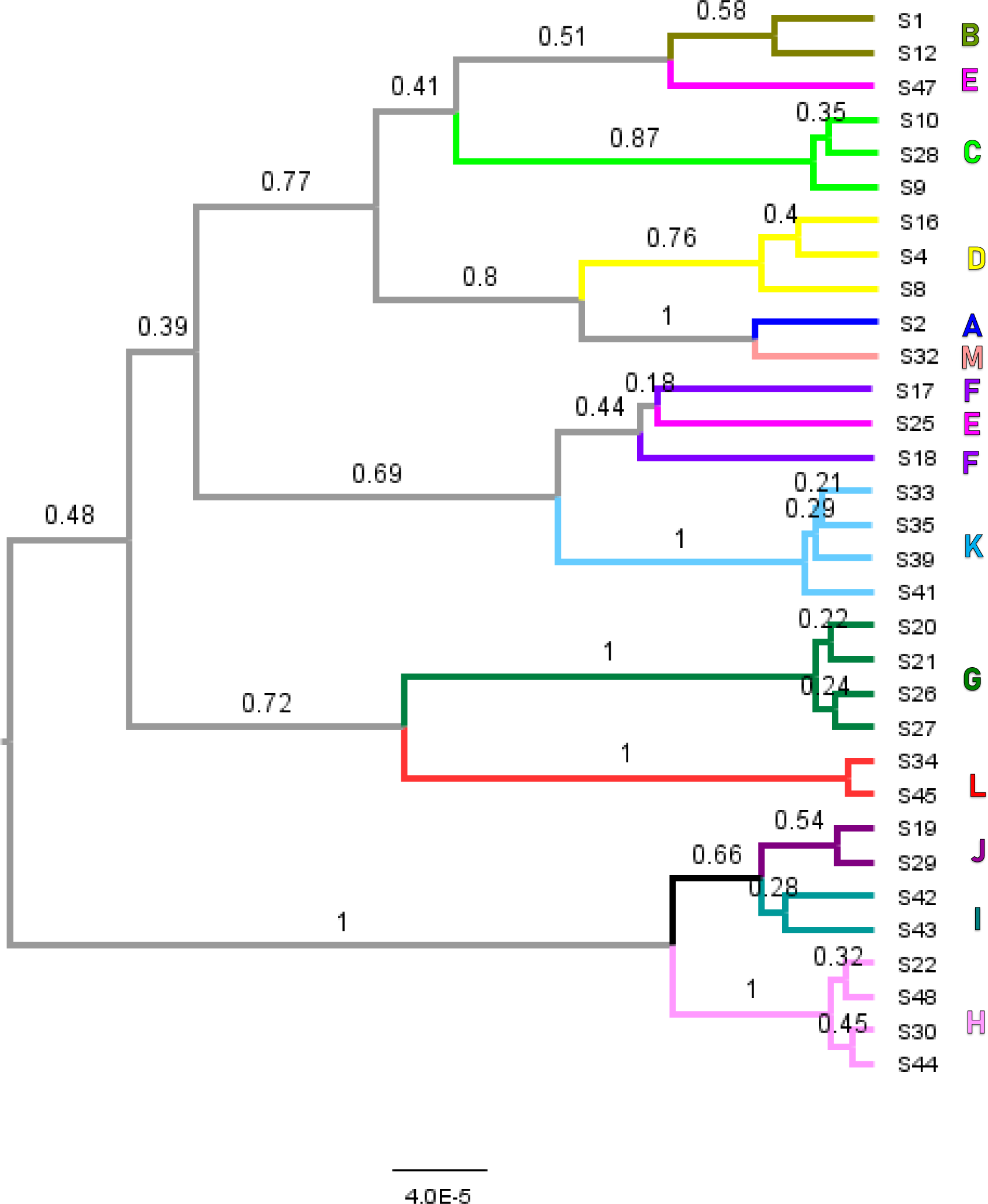
SNAPP species tree for the *Seriphium plumosum* complex and *Seriphium cinereum* estimated with 524 biallelic SNPs for 4.15 × 10^6^ generations with 1.21 × 10^6^ generations as burn-in for the best-supported species delimitation hypothesis informed by the results of STACEY, which partitions *S. plumosum* and *S. cinereum* into 32 species. Posterior probabilities are shown on branches. Tip labels and tip branch colours correspond with those in Figures 1 and 2.

### 3.5 Computation runtimes

Neither the SNAPP MCMCs based on 524 SNPs nor the STACEY analyses based on multiple alignments derived from the same 524 loci reached ESSs generally considered as sufficient (i.e. >200; BEAST 2017) for all parameters. The reduced data set with 56 randomly selected alignments showed an ESS increase rate that was more than two orders of magnitude higher compared to the 524 SNP or alignments data sets (versions 0a-1c, Table 1). The two SNAPP tree estimation analyses, together comprising 4.15 × 10^6^ generations (version 1ab, Table 1), yielded trees that were generally less well resolved than those obtained from the STACEY runs (versions 1- 3, Table 1). These two analyses also sampled less efficiently from the posterior distribution per unit time than STACEY (Table 1). The sNMF clusters G, K, and L (Fig. 1a) were recovered with posterior probability 1.00 by both STACEY (Fig. 2) and SNAPP (Fig. 3), as was the coastal HIJ clade. The clusters D, F, H, and J were also recovered in most STACEY trees (Fig 2; Supplementary Data). The STACEY runs recovered cluster I with high posterior support (Fig. 2; Supplementary Data) and usually also the clades B through K. These were not recovered by SNAPP (Fig. 3). Overall, the alternative rooting suggested by SNAPP and STACEY is perhaps the most intriguing result obtained from the comparison (Fig. 2; Fig. 3).

**Table 1:** Summary of STACEY (0a – 3) and SNAPP tree estimation (1a – 1ab) analyses. Shaded rows are replicates. See Supplementary Data for expanded table.

| Descriptor | Version | Burn-in fraction | Loci | Generations | Clusters/<br>Individuals<sup>a</sup> | ESS/day<sup>b</sup> | Resolution<sup>c</sup> |
| --- | --- | --- | --- | --- | --- | --- | --- |
| 56-alignment data set | 0a | 0.10 | 56 | 1 x 10<sup>9</sup> | 0.88 | 584.13 | 9.64 |
|  | 0b | 0.10 | 56 | 1 x 10<sup>9</sup> | 0.89 | 471.27 | 8.30 |
|  | 0c | 0.10 | 56 | 1 x 10<sup>9</sup> | 0.89 | 496.07 | 8.96 |
| Alignments unedited | 1 | 0.66 | 524 | 8.91 x 10<sup>9</sup> | 0.95 | 1.04 | 12.00 |
| Alignments with corrections | 2a | 0.71 | 524 | 5.54 x 10<sup>9</sup> | 0.88 | 2.25 | 11.00 |
|  | 2b | 0.22 | 524 | 5.72 x 10<sup>9</sup> | 0.87 | 2.91 | 10.64 |
|  | 2c | 0.90 | 524 | 4.76 x 10<sup>9</sup> | 0.96 | 1.49 | 11.00 |
|  | 2d | 0.30 | 524 | 6.07 x 10<sup>9</sup> | 0.94 | 2.49 | 10.00 |
| Identical to 2a-2d, but with all clock rates equal (=1) | 3 | 0.38 | 524 | 5.69 x 10<sup>9</sup> | 0.95 | 5.49 | 13.97 |
| SNAPP tree estimation | 1a | 0.10 | 524 | 2.066 × 10<sup>6</sup> | NA | 1.38 | 8.35 |
|  | 1b | 0.48 | 524 | 2.084 × 10<sup>6</sup> | NA | 1.46 | 7.84 |
| SNAPP tree estimation, runs 1a and 1b combined | 1ab | 0.29 | 524 | 4.15 × 10<sup>6</sup> | NA | 0.55 | 7.75 |

<sup>a</sup> Number of clusters (sensu Jones et al. 2015) divided by the number of accessions. Note that in 0a-0c, the three accessions from population 2 were coded as independent species.
<sup>b</sup> Mean effective sample size (ESS) per day calculated as [(posterior ESS + likelihood ESS)/2]/analysis runtime in days.
<sup>c</sup> Sum of posterior probability supports for clades recovered by both STACEY and SNAPP (see Supplementary Data for details).

## 4 Discussion

We present a series of analyses which demonstrate strong support for multiple species within the *S. plumosum* species complex. Several of the recovered clusters indicate a genetic basis for morphological entities recognised in a previous taxonomic scheme (Levyns 1937). However, methods differ in their degree of support for these entities, and in the relationships they recover. We compare the behaviour and performance of sNMF, STACEY and BFD, and consider the insights provided by the present work for future empirical implementations of STRUCTURE-like and MSC-based approaches for species delimitation.

### 4.1 Clusters recovered by STACEY and sNMF

A common property of the methods applied here, and indeed of most contemporary species discovery methods, is that they identify non-hierarchical clusters of objects. The sNMF method, as well as other methods based on the original STRUCTURE model (Pritchard et al. 2000), groups alleles into clusters which are not grouped further. The software SpeciesDelimitationAnalyzer, in contrast, summarizes the posterior distributions of non-hierarchical clusters recovered by STACEY where the objects clustered are minimal groups of individuals (e.g., organisms) defined *a priori*.

The clustering of alleles versus minimal clusters (Species/Minimal Clusters sensu Jones et al. 2015) is a fundamental difference, as clusters of alleles (as recognised by sNMF; Fig. 1a) may be incongruent with the clusters of organisms (as recognised by MSC-based methods; Fig. 2; Table 2) within which they reside. Individuals containing alleles from different clusters of alleles are usually interpreted as evidence of recent or past hybridization (Pritchard et al. 2000; Frichot et al. 2014). Most implementations of the MSC model, in contrast, assume no such hybridization. Instead, under the classical Wright-Fisher model, reproduction within clusters is expected to be panmictic, while reproduction between clusters is assumed to be non-existent (Hein et al. 2004). Where between- cluster hybridisation is absent, we might expect the clusters recovered by these methods (e.g., STACEY) to be equivalent to clusters recovered by admixture models (Carstens et al. 2013) provided the molecular data being used are identical. Even in the absence of hybridisation, however, incongruence between clusters could also stem from the parameterization of the MSC which, in contrast to STRUCTURE and STRUCTURE-like methods, takes into account population genealogy.

**Table 2:** Results of BFD analysis for the *Seriphium plumosum* complex and *Seriphium cinereum*. All analyses were run for 3 × 10^5^ generations.

| Species delimitation hypothesis | Number of species<sup>a</sup> | Marginal likelihood estimate | Bayes Factor | Rank |
| --- | --- | --- | --- | --- |
| Koekemoer's (2016) taxonomy | 1 | -7875.41 | 4836.26 | 7 |
| Levyns' (1937) taxonomy | 3 | -7748.81 | 4583.06 | 6 |
| sNMF1:<br>split (BC), (D), (E), (F)<br>split (I), (J) | 11 | -7406.13 ± 0.41<sup>b</sup> | 3897.70 | 5 |
| sNMF2:<br>split (BC), (D), (E), (F)<br>merge (I+J) | 10 | -7400.92 | 3887.28 | 4 |
| sNMF3:<br>merge (B+C+D+E+F)<br>split (I), (J) | 8 | -7399.89 | 3885.22 | 2 |
| sNMF4:<br>merge (B+C+D+E+F)<br>merge (I+J) | 7 | -7400.58 | 3886.60 | 3 |
| STACEY species or minimal clusters | 30 | -5457.28 | - | 1 |

<sup>a</sup> Species in *S. plumosum*.
<sup>b</sup> Mean and standard deviation for five replicate analyses.

A clear result from this study is that sNMF recovers some clusters of individuals with very little evidence of hybridisation (e.g., cluster G, Fig. 1a) which are more inclusive than the STACEY clusters. In the case of cluster G, the same four accessions were analysed, but where the sNMF analysis recovered a single cluster, the STACEY analysis recovered at least three (Fig. 2). The cluster G individuals are nevertheless unambiguously recovered as a clade by both SNAPP and STACEY (Fig. 2; Fig. 3). Similar patterns are seen repeatedly in, for example, clusters D and H-L, although some of these are ambiguously identified as either one or several clusters (D, I, J). Interestingly, within-cluster mixing is clearly more frequent among populations distributed in the western Cape Floristic Region (A through G: Fig. 1a; Supplementary Data), with populations in the eastern part of South Africa generally showing lower rates of gene pool mixing (H through L: Fig. 1a; Supplementary Data). The first-diverging lineage (Fig. 2) in the complex, population 2 (cluster A), is an exception to this general geographical trend, showing little evidence of gene pool mixing in the majority of sNMF major modes (Fig. 1a; see Supplementary Data). The overarching tendency for sNMF (Fig. 1a) to recover more inclusive clusters than STACEY (Fig. 2) provides new insights into the utility of sNMF and similar algorithms, which are generally applied at the population genetic level, for resolving higher-level lineages we typically associate with species. The smaller, less inclusive clusters supported by STACEY may be explained by the apparent deviation of our molecular data (Fig. 1a) from the Wright-Fisherian assumption of no hybridisation (Jackson et al. 2016; Leaché et al. 2019). Accommodating gene flow in the form of added model parameters should ameliorate the tendency for MSC-based methods to overestimate cluster number (Jackson et al. 2016; Leaché et al. 2019), at least so long as model violation is exclusively the result of hybridisation. Indeed, the recently developed DENIM model for species tree inference can accommodate moderate migration levels (Jones 2019a), and it is in principle possible to combine STACEY and DENIM, although this has not yet been attempted in practice. Like the “gdi” index proposed by Jackson et al. (2017), but possibly lacking in its shortcomings, DENIM offers a parameter, termed the migration decay index, which controls the rate at which migration decreases following initial divergence. Overall, a comparison of discovery MSC-based delimitation methods and sNMF using one molecular data set would be highly useful, but at present, all discovery MSC-based methods use full sequences while sNMF and other computationally efficient STRUCTURE-like methods (i.e., Snapclust and fastSTRUCTURE) are only able to compute Q matrices using SNPs.

### 4.2 Discovery versus verification

#### Species delimitation

A salient distinction between the sequence-based STACEY method (Jones et al. 2015; Jones 2017) and the SNP- or AFLP-based BFD (Leaché et al. 2014a) method of species (or population) delimitation is that, in replacing the standard *BEAST birth-death model with one that includes a spike in the node height density function near zero (the birth-death-collapse model), STACEY is able, in principle, to assess all possible assignments of accessions to species or minimal clusters. Consequently, it is able to discover such clusters without the need for an *a priori* assignment of accessions to species, hence its characterization as a “discovery” method. By contrast, BFD is a “validation” method, meaning that the user is required to formulate alternative species delimitation hypotheses *a priori*. We computed marginal likelihood estimates for seven species delimitation hypotheses (Table 2), which represent just a fraction of the full suite of possible delimitation schemes. Confining marginal likelihood estimation to a targeted set of genomically- and taxonomically-informed hypotheses may seem superficially advantageous, but there is no guarantee that the set of hypotheses thus formulated will include the correct delimitation scheme, and there is a persistent chance of the latter being omitted from the set under consideration (Grummer et al. 2014; Yang & Zhu 2018). This highlights a more general problem in model selection of choosing among multiple potentially-incorrect models (Yang & Zhu 2018). Moreover, the heavy computational burden with full MCMCs for each delimitation scheme, plus the demanding marginal likelihood estimations, which with the best current practices (Baele et al. 2012) demand a number of MCMCs themselves, render the utility of this approach questionable.

A further practical consideration for researchers carrying out analytical species delimitation is the type of the molecular marker used. We were able to retrieve and use full alignments, thereby increasing information content per locus compared to when using SNPs alone. Single nucleotide polymorphisms are typically perceived as cheaper, in terms of both cost and effort per-marker, at least when compared with Sanger sequencing (Hörandl & Appelhans 2015; Leaché & Oaks 2017). Our novel R-based haplotyping workflow, however, demonstrates that a cost/information content trade-off can be circumvented altogether with additional bioinformatic processing of short-read data, even in the absence of a reference genome. We note that the *de novo* assembly of sequences from GBS data is a capability also offered by Stacks 2.0 (Rochette et al. 2019). An important advantage of our approach is that in leveraging the location of confidently-supported SNPs to mine full sequences, the resulting sequences (like the SNPs themselves; Nosil, 2012) are concentrated in non-coding regions. This is convenient because the MSC assumes that the loci being used are neither under selection, nor pleiotropically linked to loci under selection (Felsenstein 2004). It seems reasonable to assume that data assembled in this fashion (or by Stacks 2.0) are more likely to conform to the MSC’s assumption that the alleles are neutrally evolving. The use of full sequences, as opposed to SNPs, also obviously increases the information content of the data, which should greatly enhance the resolution of MSC species trees obtained from them (Fig. 2; Fig. 3; Heled & Drummond 2010; Ogilvie et al. 2016; Jones 2019a). For example, where our STACEY analysis was informed by 3234 parsimony-informative sites including SNPs, indels and complex events in 524 sequence alignments, our BFD/SNAPP analysis was informed by only 524 biallelic SNPs, all but one SNP per locus having been removed to satisfy the assumption of marker independence in the SNAPP tree estimation step (Bryant et al. 2012).

The reduced 56-alignment STACEY analysis achieved resolution similar to the 524-SNP SNAPP tree, but at a convergence rate more than two orders of magnitude faster (Table 1). In light of the apparent need for MSC methods to include more parameters, further studies on the minimum data sizes required to achieve model stationarity appear warranted.

#### Computation speed

The BFD method was originally presented as a tree-based computationally efficient means of comparing the marginal likelihood estimates of alternative species delimitation hypotheses (including non-nested schemes) using genome-scale data (Leaché et al. 2014a). We expected BFD to emerge as more computationally efficient than STACEY, both because it uses SNPs over full sequences (see Andermann et al. 2019) and because we only considered a handful of species delimitation hypotheses (Table 2). Contrary to our expectations, the SNAPP tree estimation step in BFD run for 3 × 10^5^ generations produced a poorly-resolved tree (Supplementary Data), and even when combining two substantially extended independent SNAPP tree estimation analyses, tree resolution and ESS achieved for the posterior and likelihood per unit time were poorer (Fig. 3; Table 1) than STACEY (Fig. 2; Table 1). These results suggest that STACEY is preferred when using a moderately large number of loci and accessions, both for its relative computational efficiency for phylogeny estimation, and because it exhaustively considers every possible species delimitation scheme. Considering the number of phased allelic individuals in our data set (n = 72) corresponding to 36 tips in the species tree, and the large number of delimitation schemes possible, we consider the computation time of STACEY remarkably low. A further shortcoming of BFD in relation to STACEY is that SNAPP removes missing data non-randomly such that more species-rich schemes in BFD are informed by fewer loci (Supplementary Data). The SNAPP trees associated with alternative species delimitation hypotheses therefore cannot be considered directly comparable. Having made these remarks, however, we note that the number of accessions we used far exceeds that for which BFD was originally intended (Leaché et al. 2014a), and acknowledge its computational viability so long as the number of accessions remains small.

### 4.3 Species as clusters versus species as clades

Current species delimitation methods based on the MSC identify the terminal branches of the MSC tree as the extant species (and the internal branches as ancestral species, see Degnan and Rosenberg 2009). However, the parameterization available may be insufficient to adequately identify biologically relevant clusters. Although extensive simulation studies have been performed assessing the MSC’s robustness to gene flow (Zhang et al. 2011; Leaché et al., 2014b; Greunstaeudl et al. 2016; Barley et al. 2018), there is an urgent need to incorporate additional parameters tin an effort to improve the biological realism with which we model gene tree discord beyond discord caused by the incomplete sorting of alleles and (low levels of) hybridisation.

An even more important consideration is whether species should be conceptualized as a category distinct from clades and populations (see de Queiroz 2020). The MSC provides a framework for conceptualizing species as the branches of the species tree, where current implementations are based on Wright-Fisher assumptions, and thus applicable to identify such populations. If species are conceptualized as clades, species delimitation is merely a problem of ranking. Conceptually, however, clades and non-hierarchical clusters are fundamentally different things. Clades are nested in one another in trees, and the inclusiveness for species (or other taxonomic ranks) is, in principle, arbitrary, but may be guided by auxiliary criteria (Dayrat 2005; Padial et al. 2009). On the other hand, it appears to be theoretically possible to parameterize species as non-hierarchical clusters of individuals under the MSC, with some individuals having some alleles descended from other clusters in the past (i.e., migration). This is a major question for biologists to consider, because in the latter case, species may be viewed as a category as fundamental and non-arbitrary as the population or clade categories. In the former case, delimitation of species will continue to lack an explicit and parameterized model through which biologists can view the fit of their data.

In conclusion, our results show that when analysing 500+ putatively independent loci for the S. plumosum complex, the number of species identified by MSC methods approaches the number of individuals sampled. The STRUCTURE-like approach sNMF identifies fewer clusters, and those with no or little admixed individuals tend to be congruent with clades retrieved by MSC methods, even when the latter otherwise show questionable mixing. Convergence rates for the MSC method STACEY increase by more than two orders of magnitude when reducing the number of loci by an order of magnitude, while still recovering strongly supported clades congruent with “clean” sNMF clusters, and with only a slight reduction in overall resolution. This suggests that future development of more realistically parameterized models should focus on few, well-curated and informative loci rather than “whole-genome” data sets.

## ACKNOWLEDGEMENTS

We thank the University of Cape Town’s ICTS High Performance Computing team (hpc.uct.ac.za), the CIPRES Science Gateway (phylo.org), UNINETTSigma2, the National Infrastructure for High Performance Computing and Data Storage in Norway (sigma2.no), and Linköping University’s National Supercomputer Centre (nsc.liu.se) for access to computational resources on “Tetralith”.

We thank Mr Terry Trinder-Smith and Miss Charlene Christians for their generous assistance at the Bolus Herbarium. We also thank John Burrows, Sandie Burrows, and Barbara Turpin of the Buffelskloof Private Nature Reserve for their assistance with field work. Professor Adam Leaché and Dr Graham Jones are thanked for helpful advice relating to model setup.

## FUNDING

This work was supported by a grant from the South African National Research Foundation, grant number 105976.

## SUPPLEMENTARY MATERIAL

Genotyping-by-sequencing data in binary alignment mapping (BAM) format and the associated *de novo*-assembled genome have been deposited in the NCBI short-read archive: BioProject PRJNA705849. Annotated R code for the *de novo* assembly of seqeunce alignments from short-read data for diploid organisms, and for large-scale STACEY XML assembly is available on GitHub: https://github.com/zaynabthebotanist. Processed genomic data files are available from the Dryad Digital Repository: https://doi.org/10.5061/dryad.sj3tx964m. Phylogenetic trees have been deposited in TreeBASE: XXX.

## Notes

https://doi.org/10.5061/dryad.sj3tx964m

https://www.ncbi.nlm.nih.gov/bioproject/PRJNA705849/

### Competing Interest Statement

The authors have declared no competing interest.

